## Supplementary Materials for "Maturational networks of human fetal brain activity reveal emerging connectivity patterns prior to ex-utero exposure"

### Supplementary Materials to: “Maturational networks of fetal brain activity”

#### *Supplementary Figures*

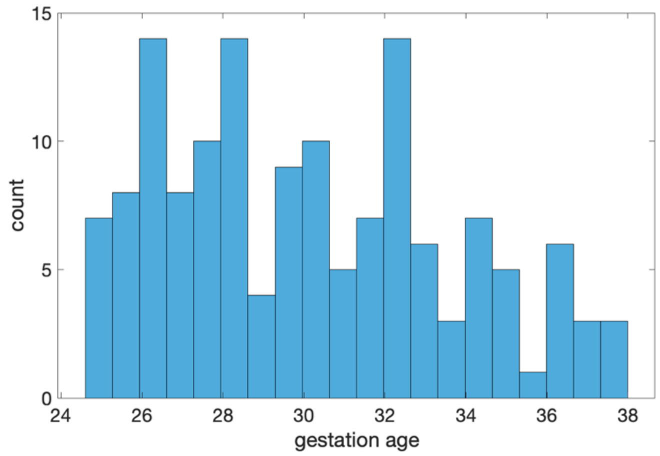

*Fig. S 1. Distribution of gestation ages in weeks in the studied sample.*

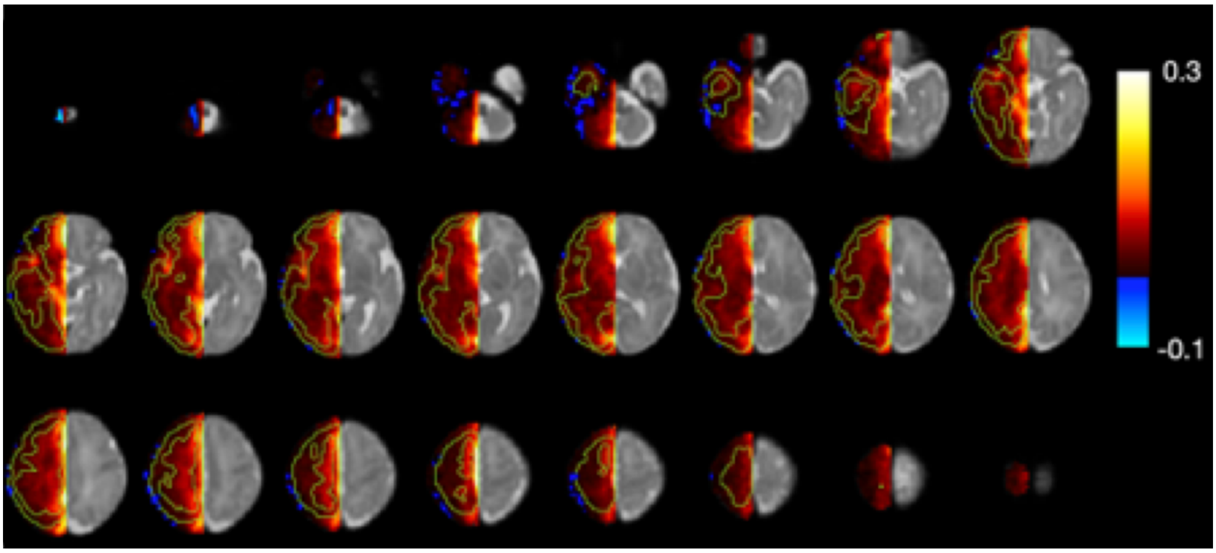

*Fig. S 2. Maps of group-average correlation between homologous voxels in the two hemispheres. The analysis is performed in a symmetrical template space. Note that for this analysis, effects of spatial and anatomical proximity on the strength of correlation and its age-related change are dissociated; in other words, a pair of homologous voxels near the medial wall, despite their spatial proximity, is anatomically as distant on average as a pair of laterally located regions.*

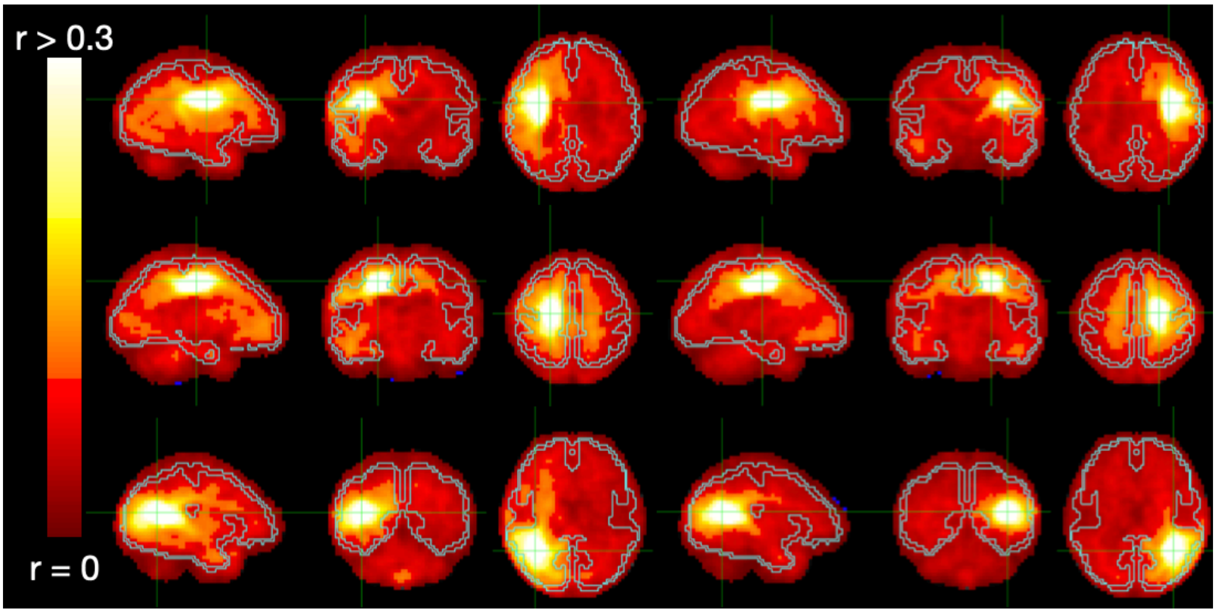

*Fig. S 3. Seed-to-brain maps of group-average correlations for the white matter seeds. Location of a seed coincides with the center of the crosshair. Examples of 6 seeds are shown, 3 per each hemisphere. The presence of distance-dependent gradient is evident. Some differences can be observed qualitatively between the white and grey matter (as shown in Fig. 2a); the white matter tends to show a broader spread of increased and spatially diffuse values away from the location of a seed, partially mirroring the shape of white matter "skeleton".*

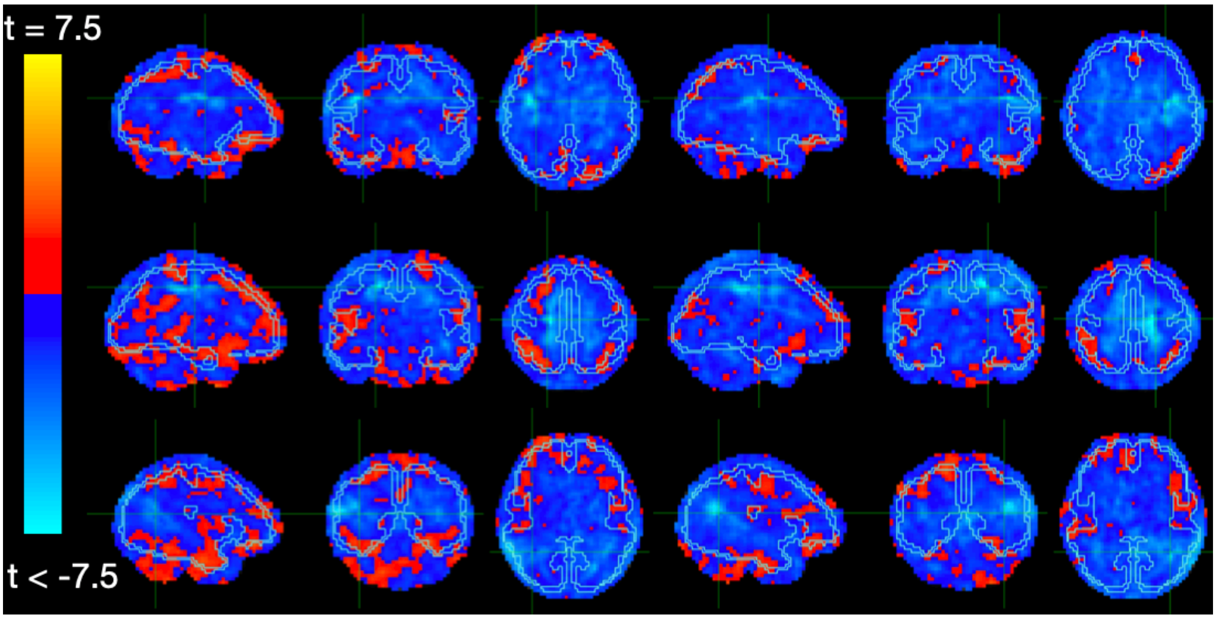

*Fig. S 4. Seed-to-brain maps of age effect (t-value map) on the strength of correlations for the white matter seeds. Location of a seed coincides with a center of the crosshair. Examples of 6 seeds are shown, 3 per each hemisphere. The increased negative values at the location of the seed (centre of the crosshair) are present but less conspicuous than in the grey matter maps. Note sparseness and weakness of the positive associations (i.e., the increase of seed connectivity with age), comparing with the grey matter seeds in Fig. 2b.*

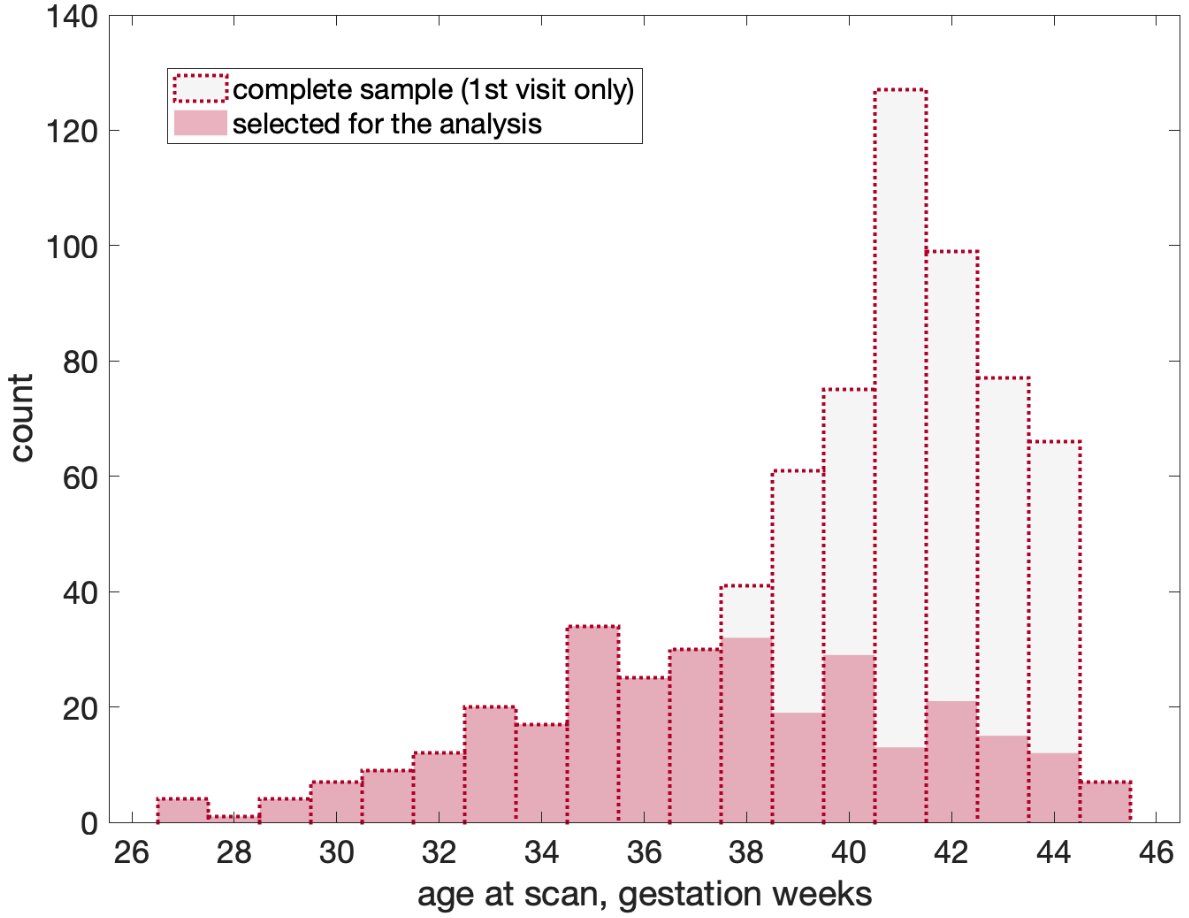

*Fig. S 5. Neonatal sample used in the study, compared to the dHCP neonatal cohort.*

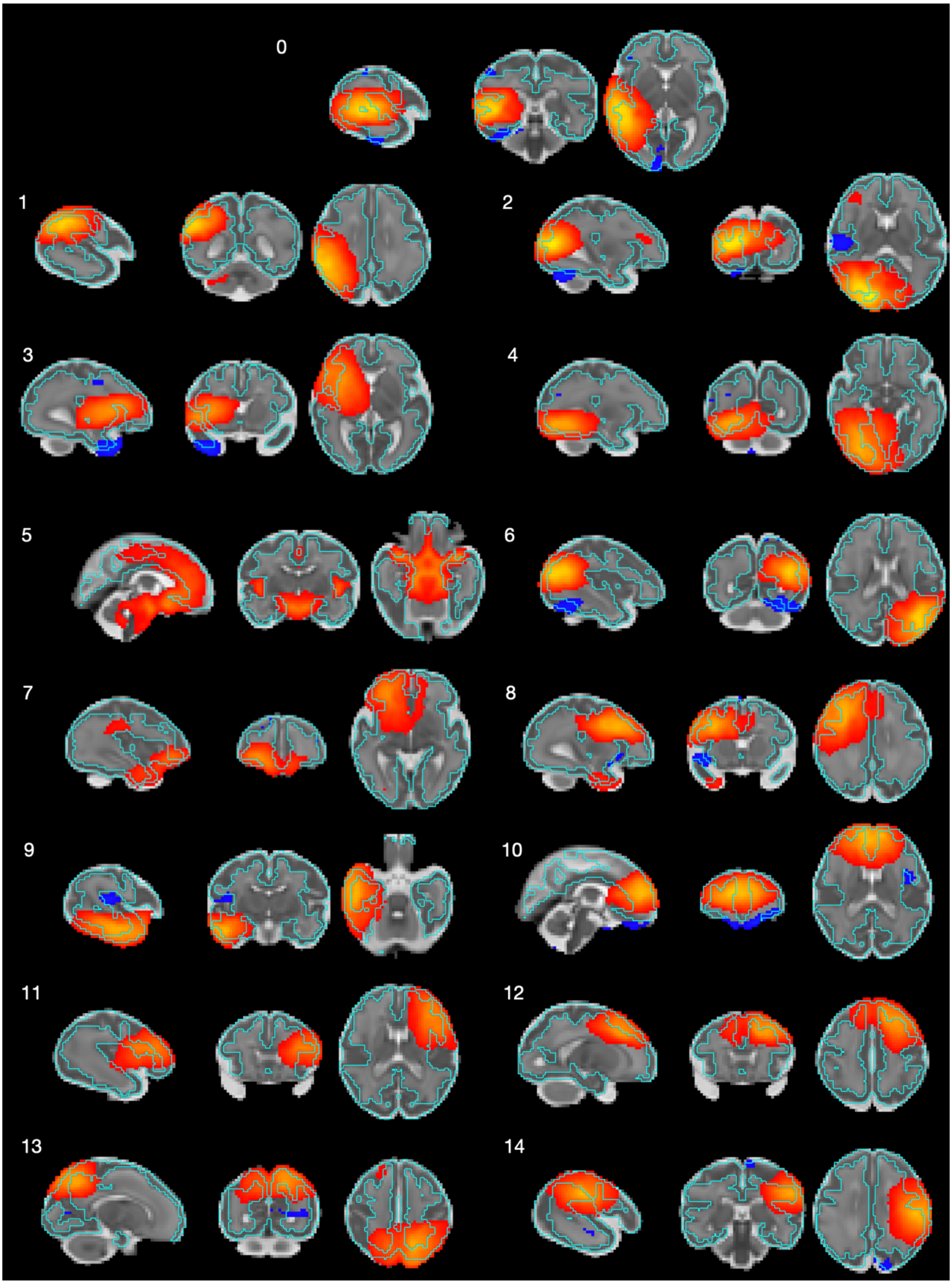

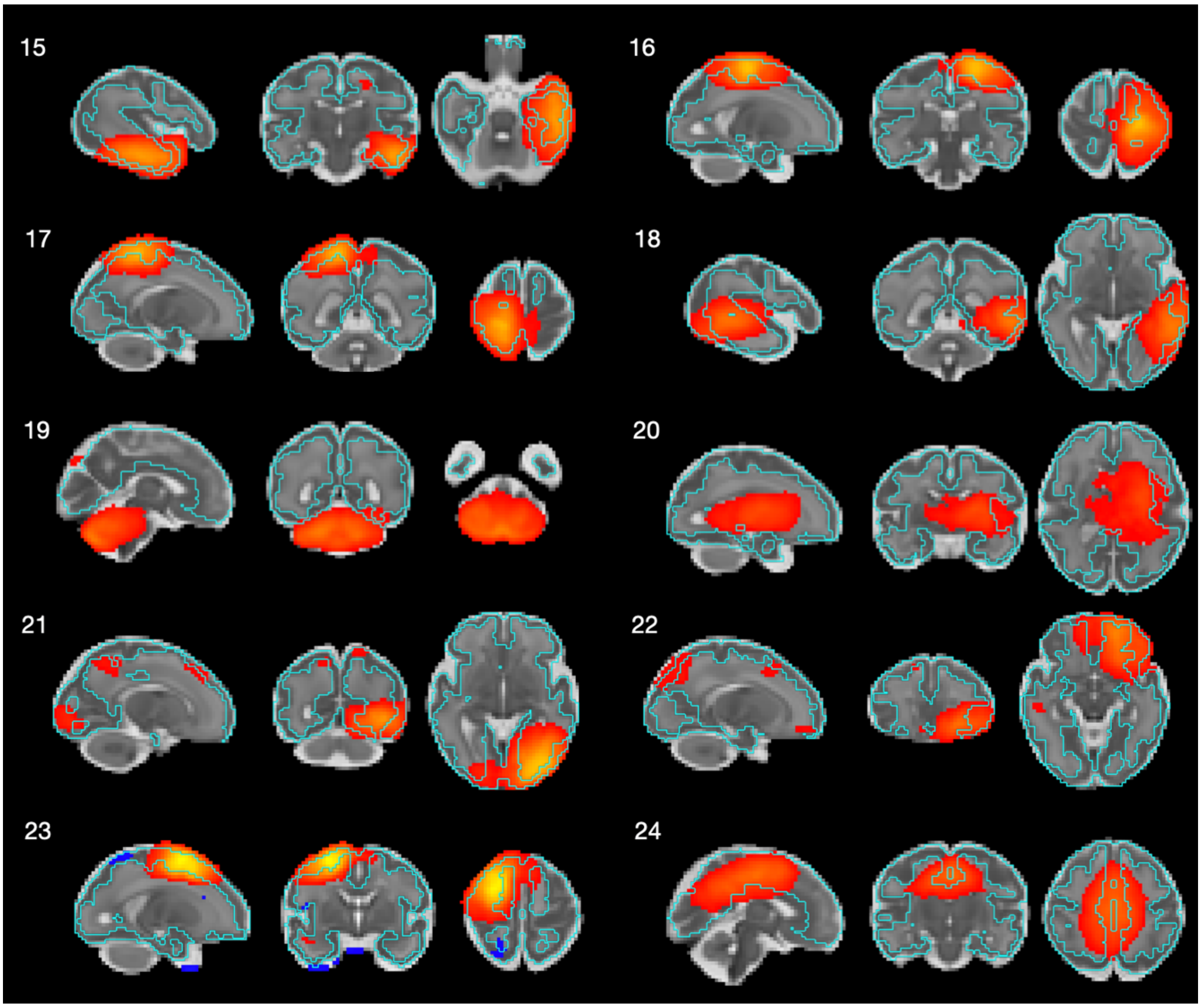

*Fig. S 6. Group ICA maps shown with overlaying cortical ribbon boundaries (shown in light blue).*

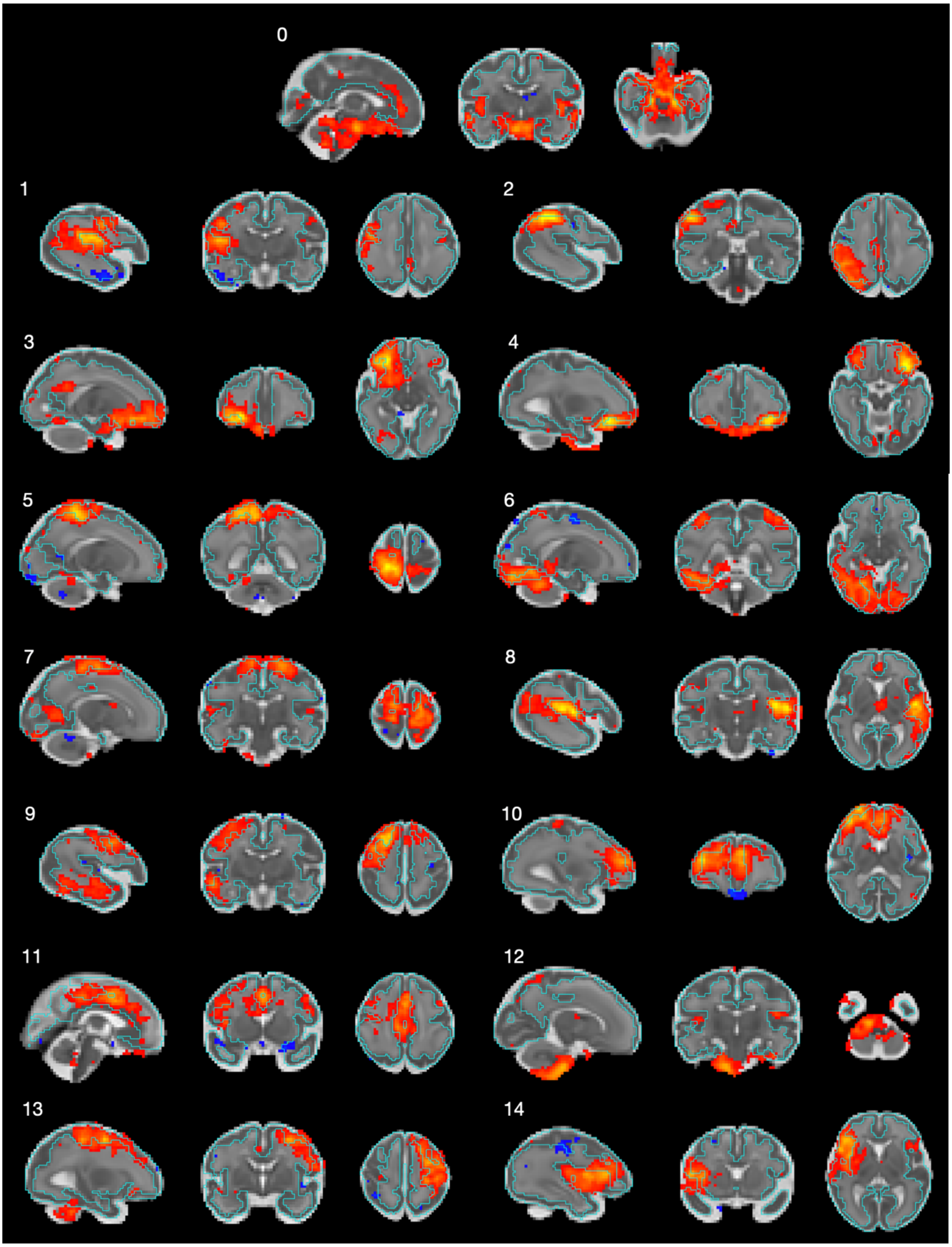

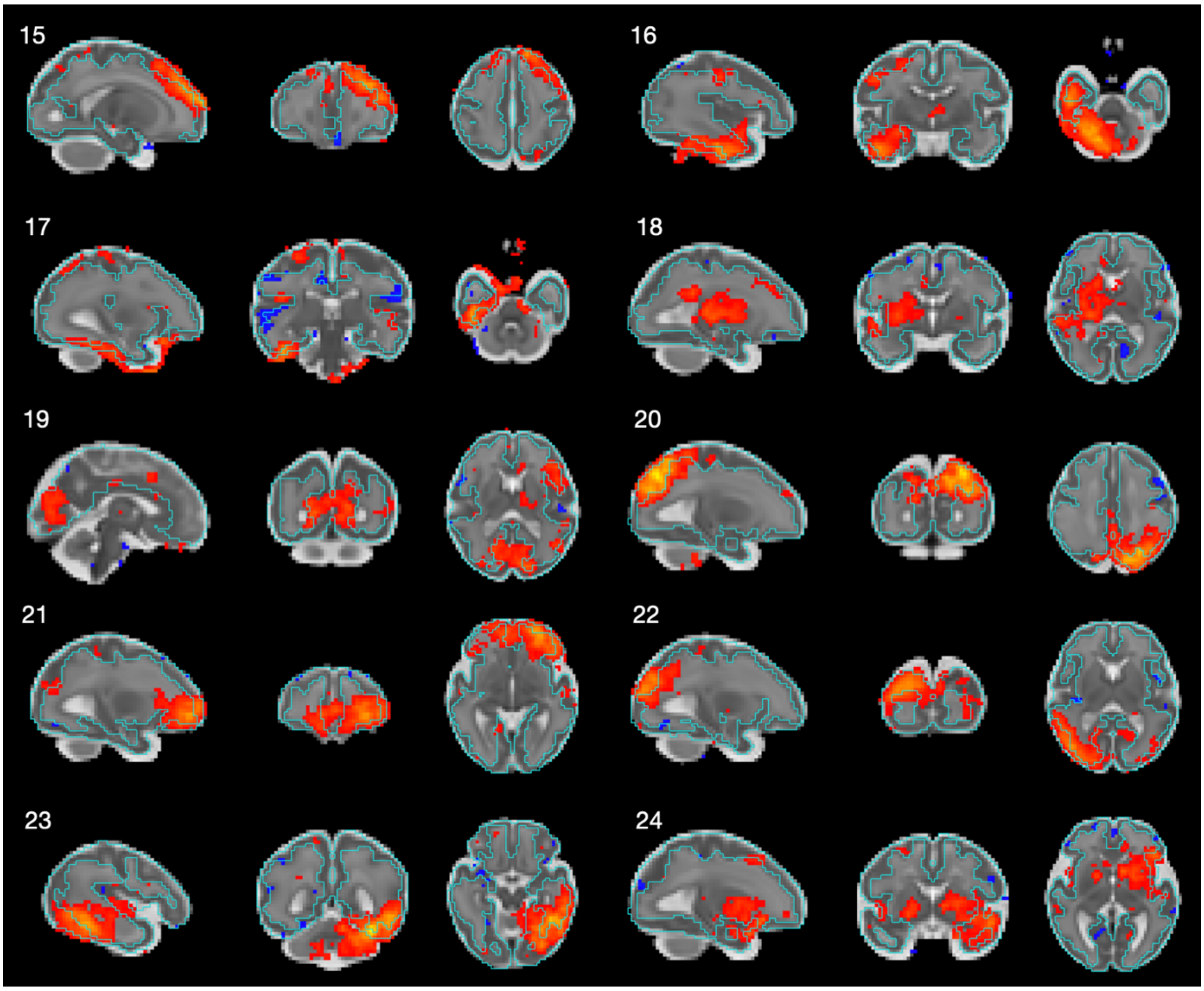

*Fig. S 7. Maturational networks shown with overlaying cortical ribbon boundaries (shown in light blue)*

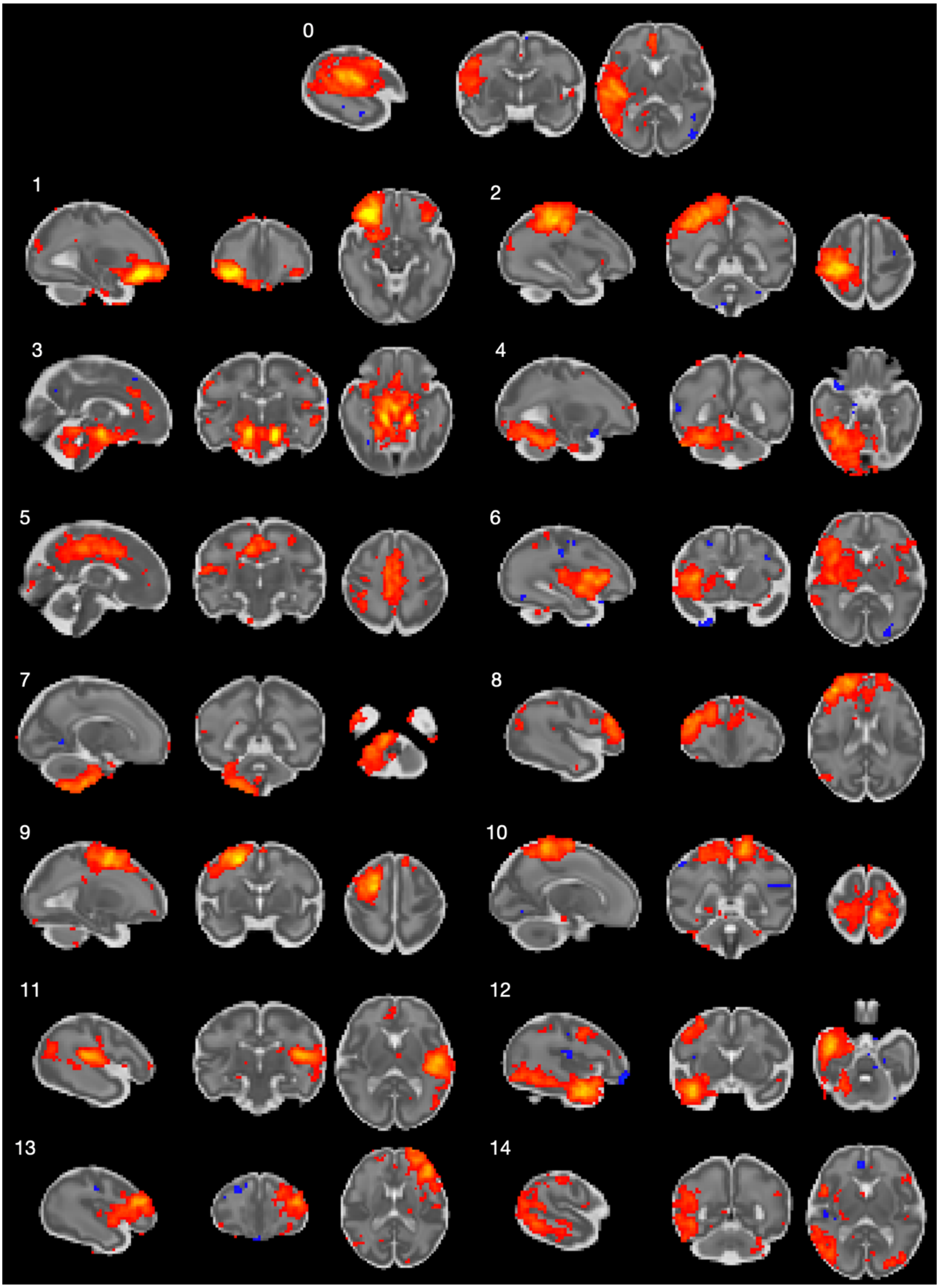

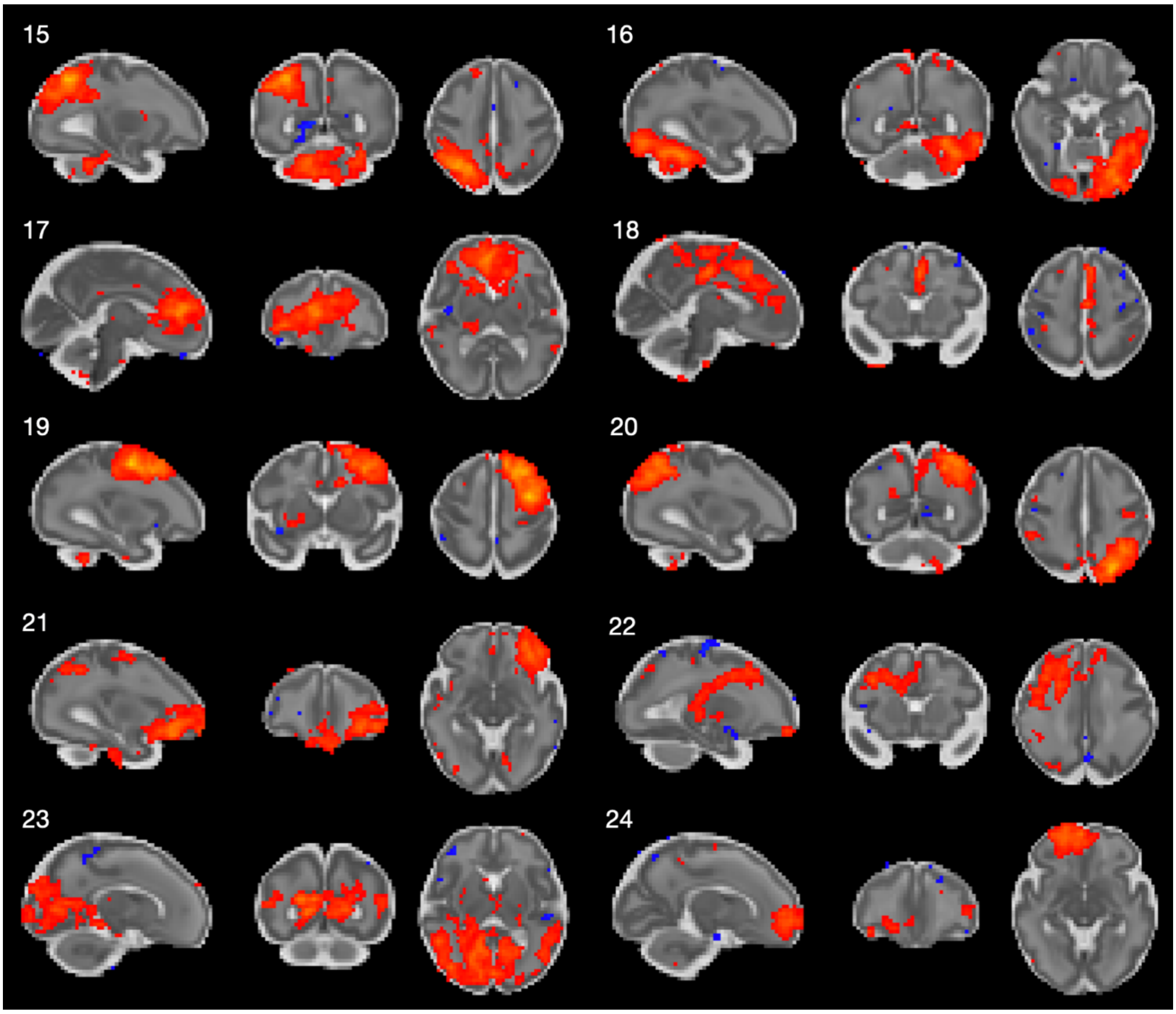

*Fig. S 8. Maturational networks – split-half sample 1*

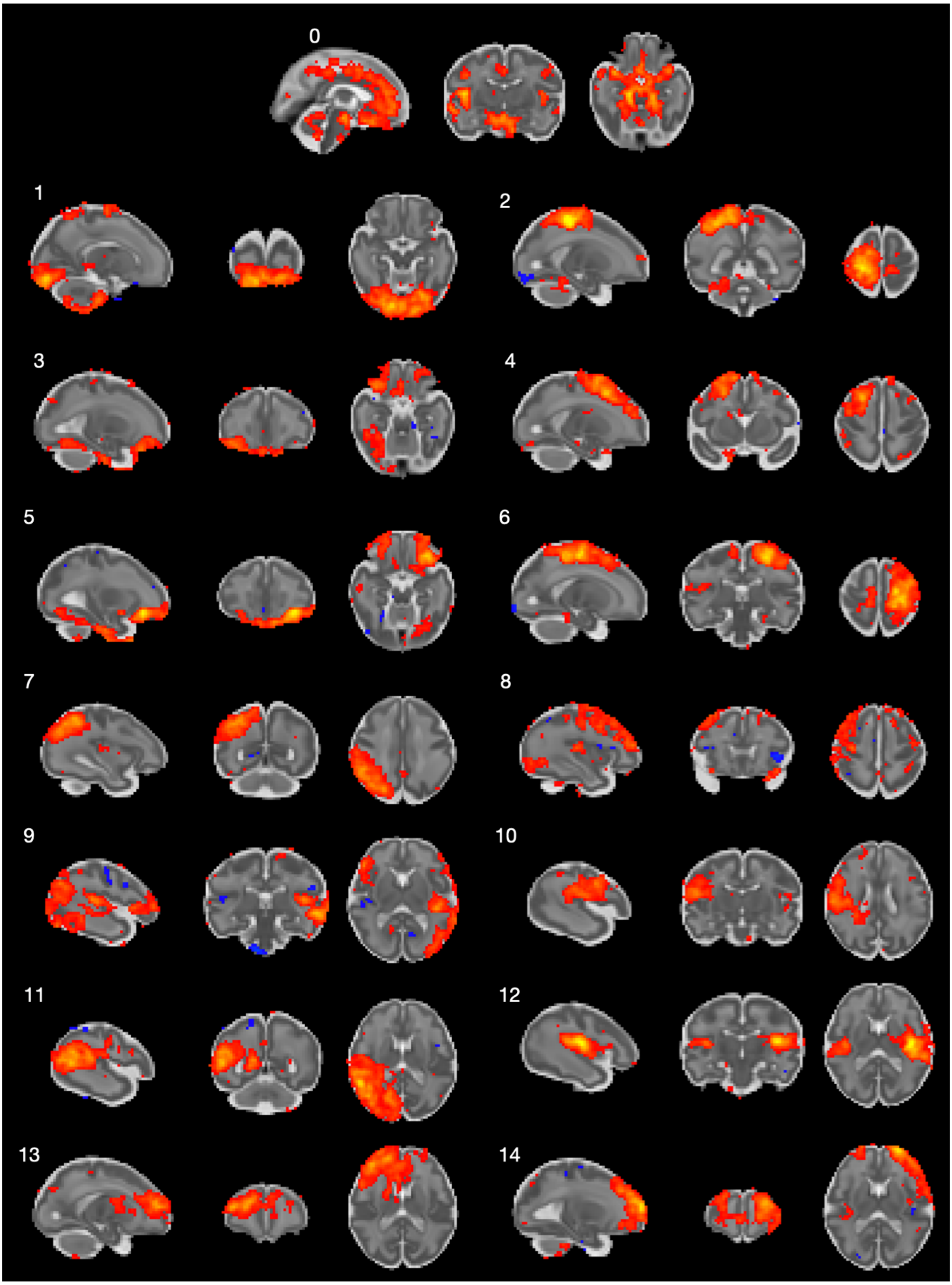

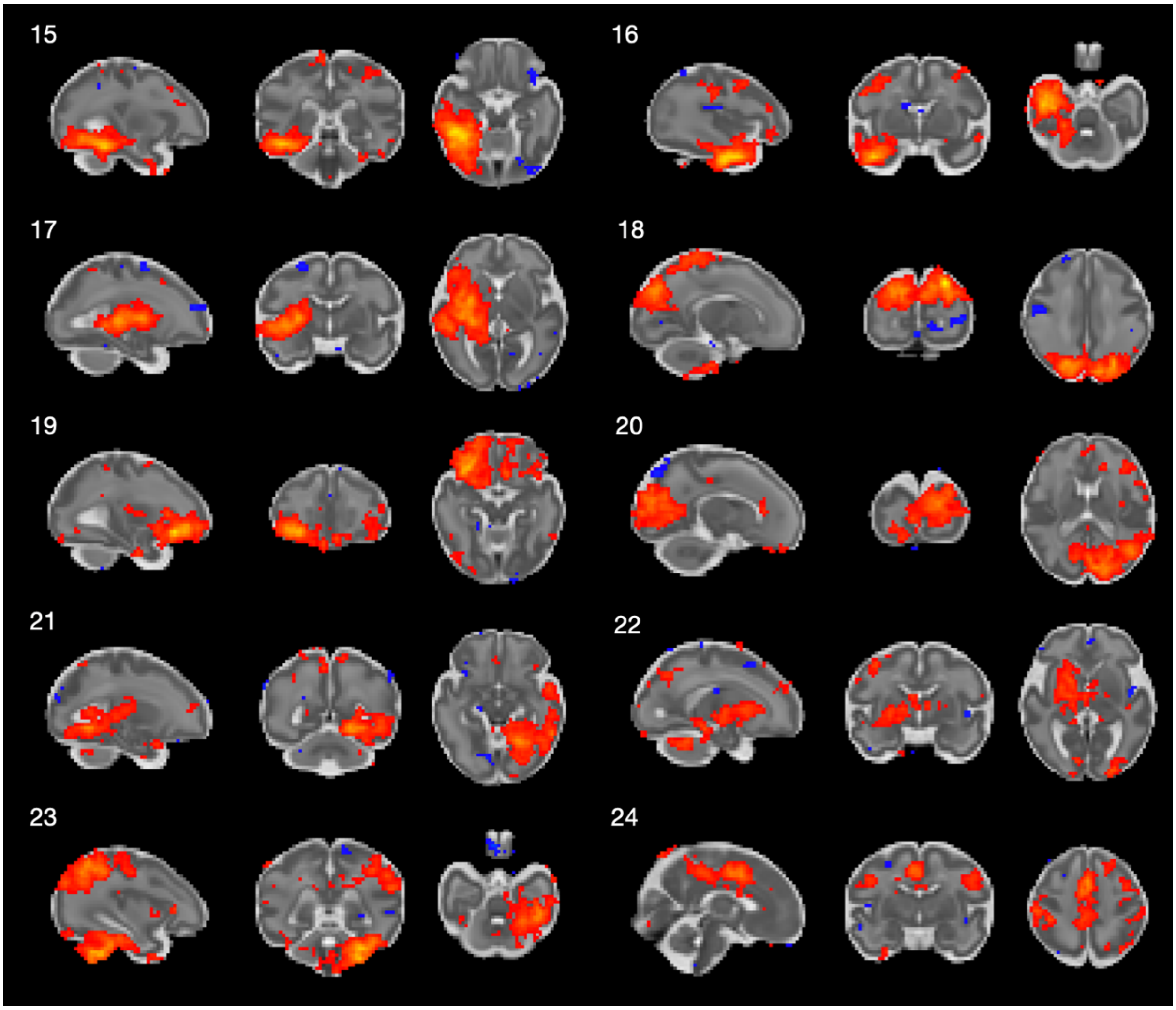

*Fig. S 9. Maturational networks – split-half sample 2*

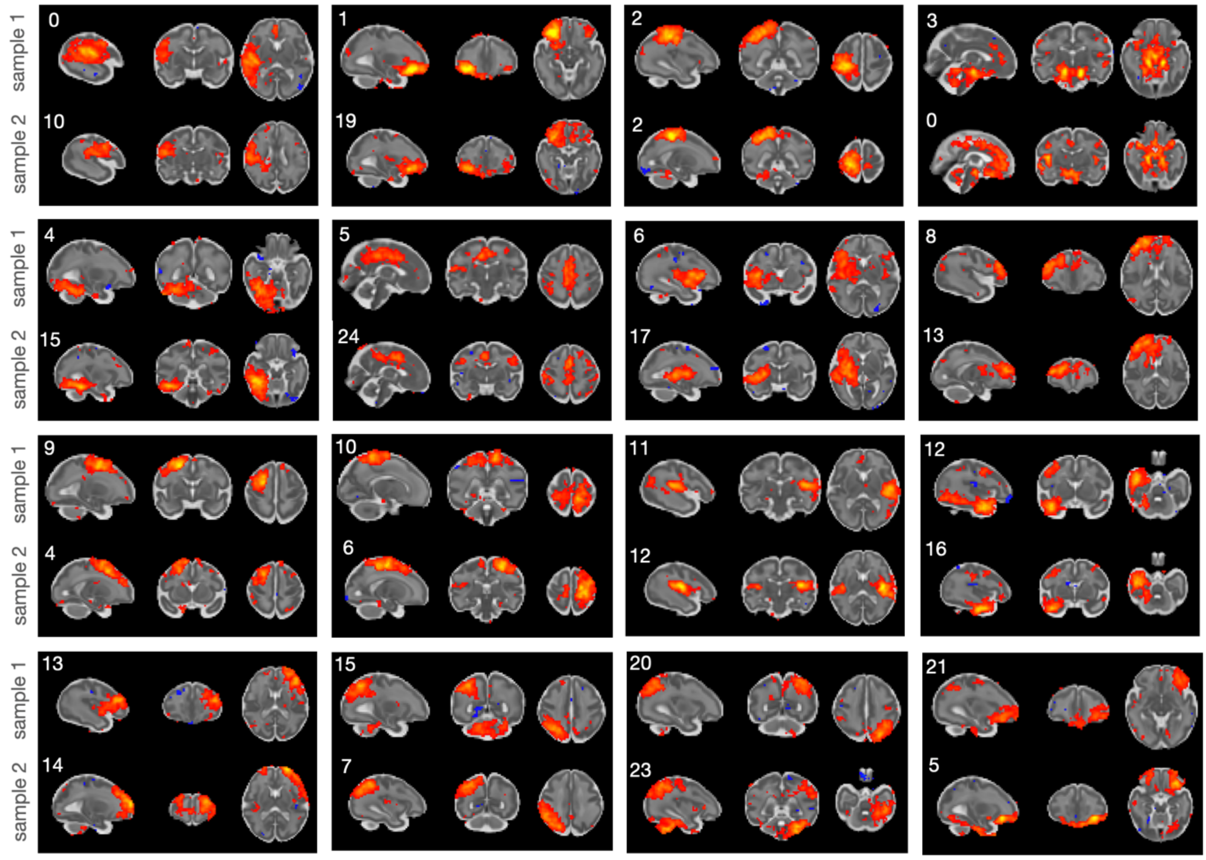

*Fig. S 10. Paired components from split-half sample 1 (top) and split-half sample 2 (bottom) based on their spatial similarity*

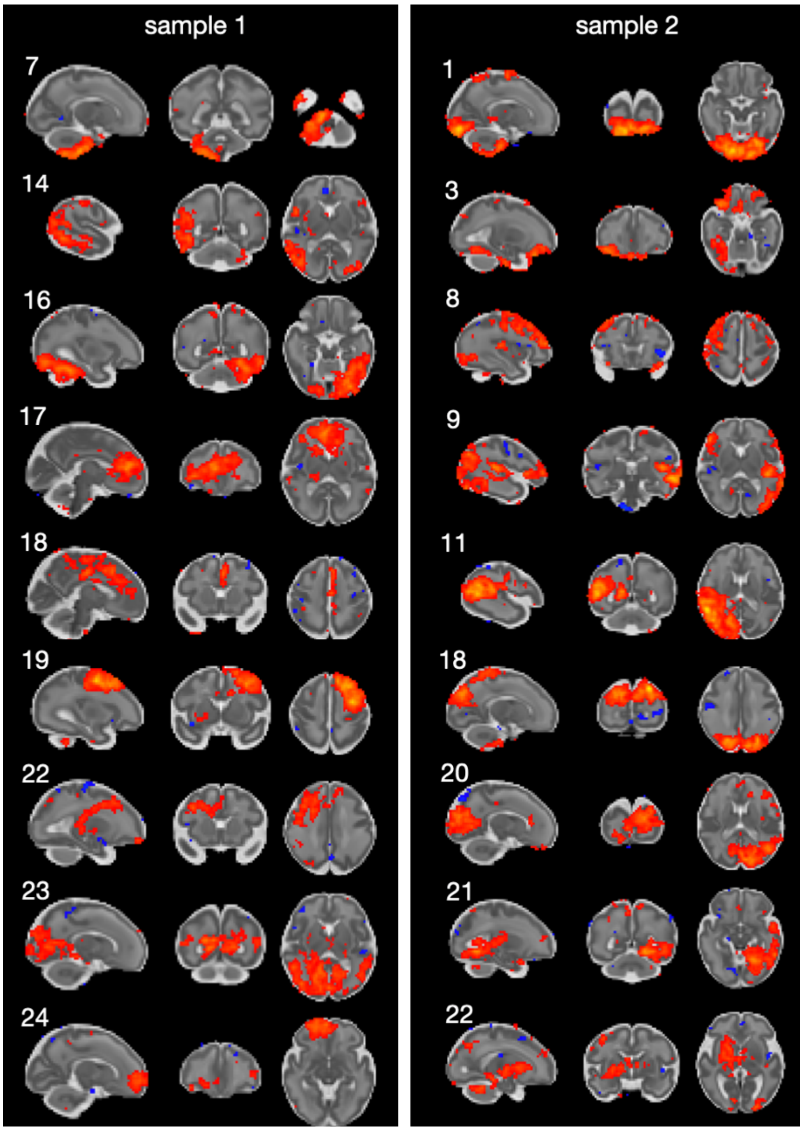

*Fig. S 11. Non-paired components from split-half samples 1 and 2.*

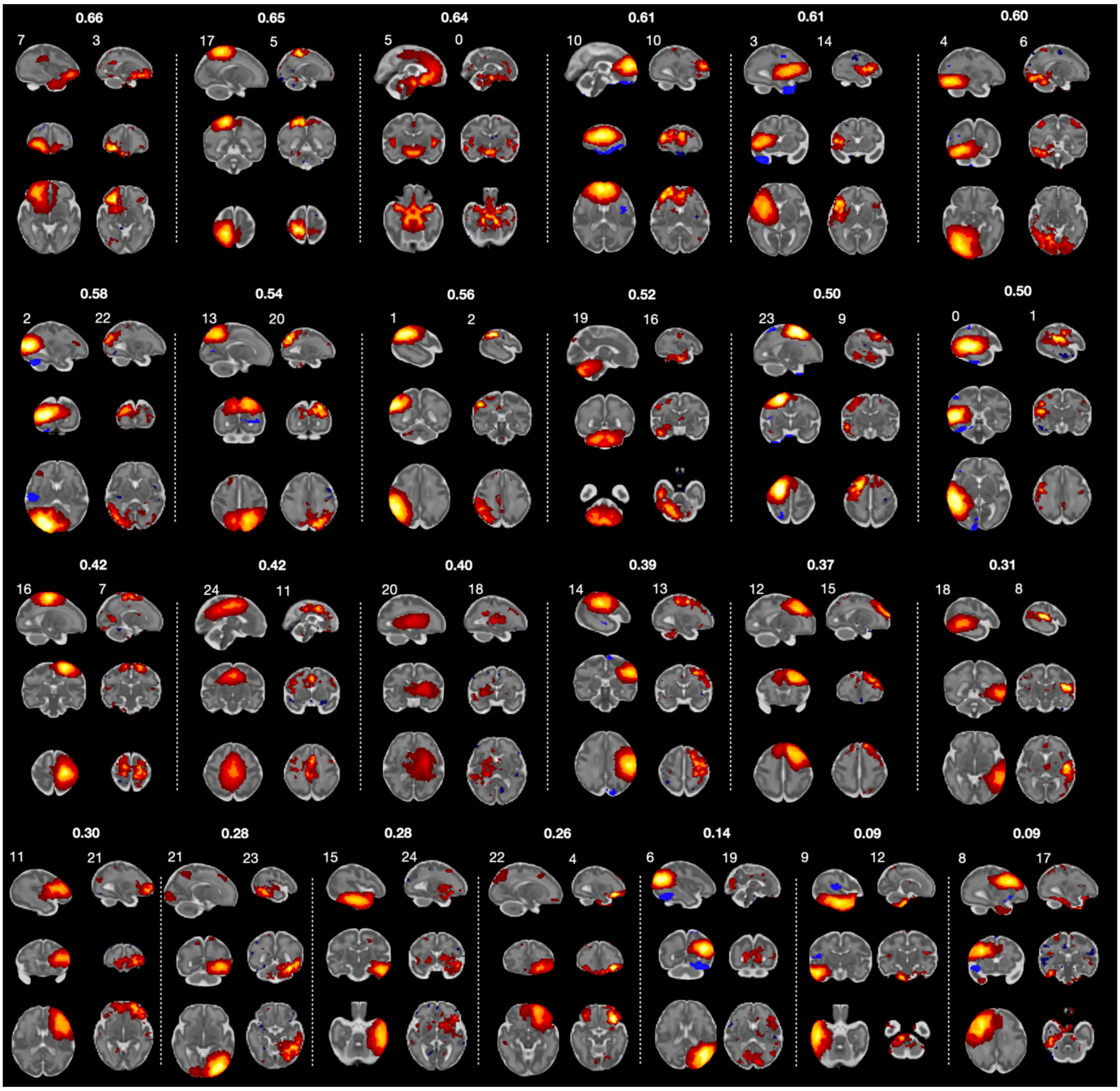

*Fig. S 12. Group-ICA (left in a pair) and matnets (right in a pair) paired using Hungarian algorithm based in their similarity. The pairs arranged in an ordered manner, starting from fairly well matched (= high spatial correlation) to unmatched (= low spatial correlation).*

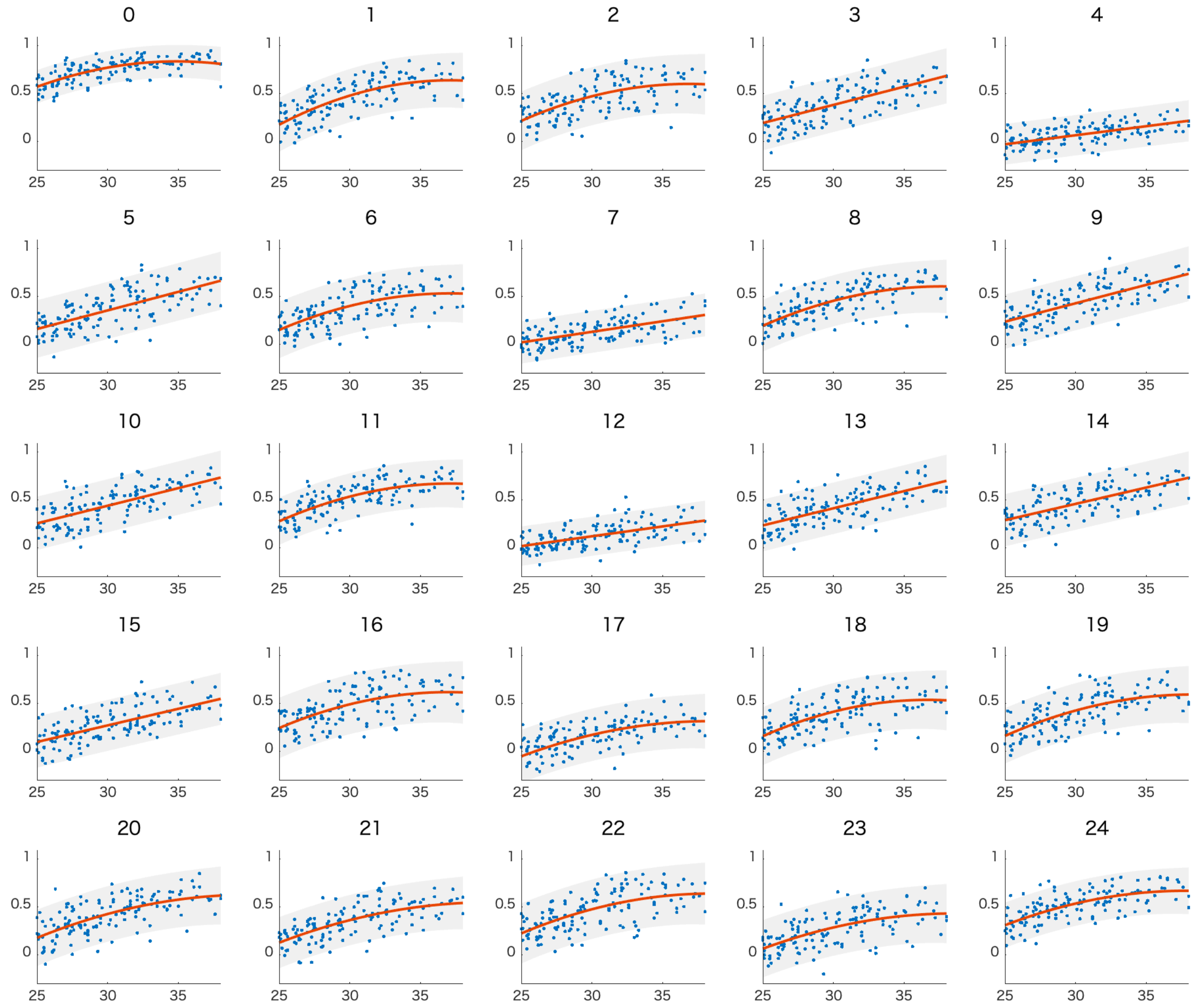

*Fig. S 13. Temporal correlations between time courses of maturational components and their complementary maps. All maps were thresholded at z >5 in order to reduce a degree of potential spatial overlap between pairs. The time courses were computed as weighted averages of the above-threshold voxels.*

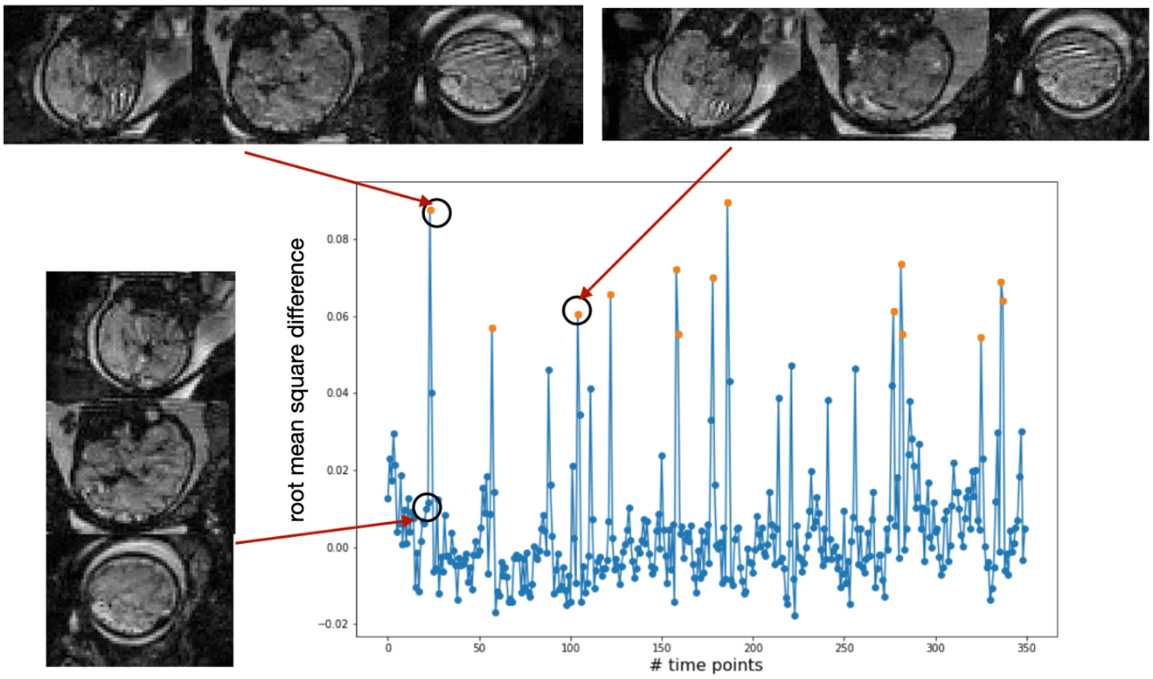

*Fig. S 14. Volume censoring. Orange dots highlight censored volumes (root mean square intensity difference from median normalised with respect to the grand average median > .05)*

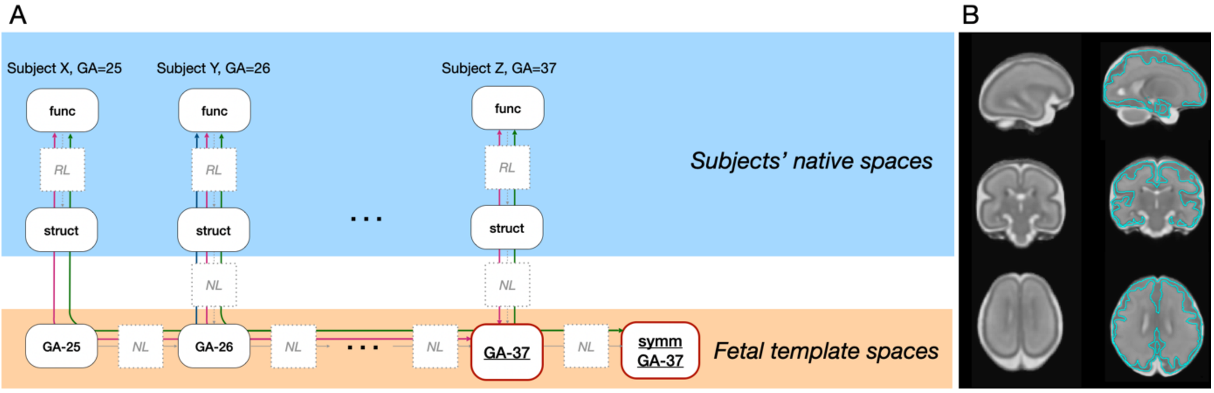

*Fig. S 15. Registration to spaces for group-level analyses. A). Registration pipeline. Func - functional data; struct – anatomical scan; GA-25 etc. – brain template corresponding to 25 weeks of gestation etc.; symm – symmetrical template space; RL – rigid linear registration; NL – non-linear diffeomorphic registration. B) Example of the registration of the youngest template (GA-25) to the group space (GA-37). Left – original GA-25 template. Right – the GA-25 template warped into the GA-37 template space. Despite that gyrification is not well developed at the age of 25 gestation weeks, concatenation of the sequential warps allows for an accurate deformation, as demonstrated by the cortical outline of GA-37 template, overlaying the warped image.*

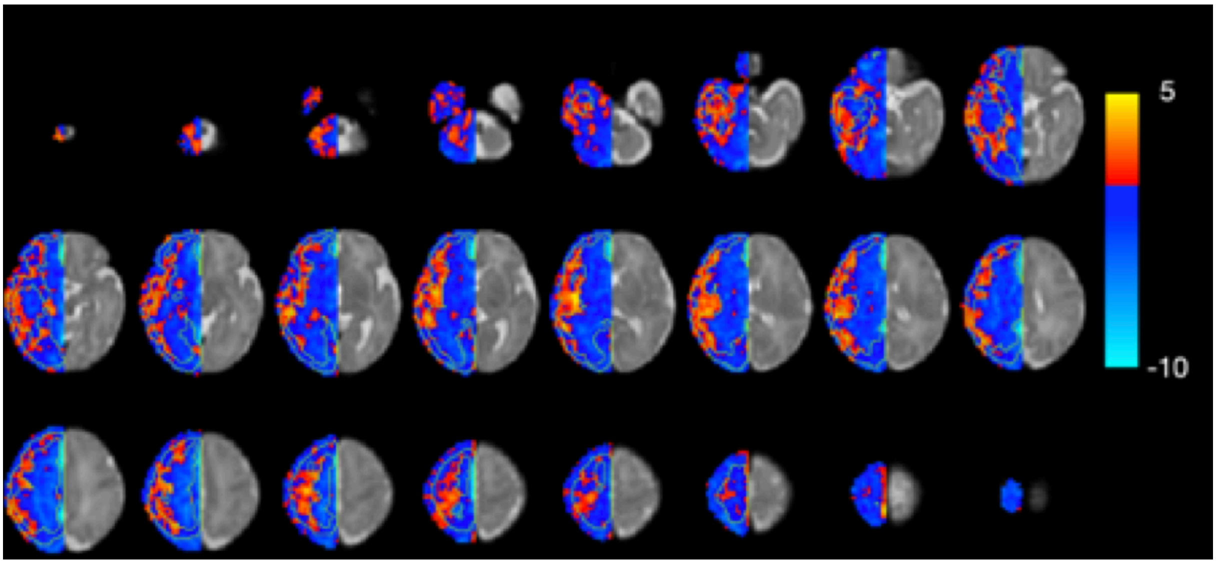

*Fig. S 16. Age-related changes in connectivity between homologous voxels in the two hemispheres. The map was used to define the seeds for the univariate seed-to-brain analysis. Grey matter seeds were generated by thresholding the map positively at z >3. White matter seeds were generated by thresholding the maps negatively, and disregarding areas near the medial wall. Three clusters in the proximity of the grey matter seeds were manually adjusted to match their size to enable qualitative comparisons between the univariate signal properties of the two tissue classes.*
